# Chronic alcohol drinking persistently suppresses thalamostriatal excitation of cholinergic neurons to impair cognitive flexibility

**DOI:** 10.1101/2021.10.18.464867

**Authors:** Tengfei Ma, Zhenbo Huang, Xueyi Xie, Yifeng Cheng, Xiaowen Zhuang, Matthew Childs, Himanshu Gangal, Xuehua Wang, Laura Smith, Rachel Smith, Yubin Zhou, Jun Wang

**Affiliations:** Department of Neuroscience and Experimental Therapeutics, College of Medicine, Texas A&M University Health Science Center, Bryan, Texas, USA; Department of Psychology, Texas A&M University, College Station, Texas, USA; Institute of Biosciences and Technology, Department of Translational Medical Sciences, College of Medicine, Texas A&M University, Houston, Texas, USA

**Keywords:** alcohol, cholinergic interneuron, striatum, glutamatergic transmission, thalamus, behavioral flexibility

## Abstract

Exposure to addictive substances impairs flexible decision-making. Cognitive flexibility is mediated by striatal cholinergic interneurons (CINs). However, how chronic alcohol drinking alters cognitive flexibility through CINs remains unclear. Here, we report that chronic alcohol consumption and withdrawal impaired reversal of instrumental learning. Chronic alcohol consumption and withdrawal also caused a long-lasting (21 d) reduction of excitatory thalamic inputs onto CINs and reduced pause response of CINs in the dorsomedial striatum (DMS). CINs are known to inhibit glutamatergic transmission in dopamine D1 receptor-expressing medium spiny neurons (D1-MSNs) but facilitate this transmission in D2-MSNs, which may contribute to flexible behavior. We discovered that chronic alcohol drinking impaired CIN-mediated inhibition in D1-MSNs and facilitation in D2-MSNs. Importantly, *in vivo* optogenetic induction of long-term potentiation of thalamostriatal transmission in DMS CINs rescued alcohol-induced reversal learning deficits. These results demonstrate that chronic alcohol drinking reduces thalamic excitation of DMS CINs, compromising their regulation of glutamatergic transmission in MSNs, which may contribute to alcohol-induced impairment of cognitive flexibility. These findings provide a neural mechanism underlying inflexible drinking in alcohol use disorder.

## Introduction

Alcohol use disorder is a chronic brain disorder characterized by an inability to stop drinking despite the resultant adverse consequences (1, 2). This inability is associated with impaired flexibility in decision-making, which contributes to compulsive alcohol use (1–5). Increasing evidence suggests that the dorsomedial striatum (DMS) is involved in cognitive flexibility (6–11). Understanding whether and how chronic alcohol consumption affects striatum-mediated cognitive flexibility will provide therapeutic strategies to treat alcohol addiction.

In the DMS, cholinergic interneurons (CINs) are the major source of acetylcholine and contribute to cognitive flexibility in response to salient stimuli (12–14). CINs play an essential role in modulating striatal circuit activity, thereby regulating output from the striatum (15–17). The medium spiny neurons (MSNs), which express either dopamine D1 receptors (D1R) or D2Rs, are the principal striatal projection neurons. D1-MSNs and D2-MSNs play different roles in motor control and goal-directed behavior (18–24). Accumulating evidence demonstrates that the characteristic burst-pause firing of CINs regulates MSN activity; this firing pattern is triggered by excitatory inputs from the thalamus, which is a critical modulator of striatal activity (14, 17, 25). MSN regulation by CINs is mediated by the actions of acetylcholine on pre- and postsynaptic muscarinic receptors. For example, burst-associated transient acetylcholine release produces a muscarinic M2/M4 receptor-mediated reduction in glutamate release at corticostriatal terminals on both D1- and D2-MSNs (17, 26). The more prolonged effects of acetylcholine on postsynaptic excitability during the “pause window” are mediated by the preferential activation of muscarinic M1 receptors on D2-MSNs, but not D1-MSNs. These studies demonstrated that CIN burst-pause firing following thalamic activation is crucial for the functional modulation of striatal MSNs. Since striatal D1- and D2-MSNs respectively give rise to the direct (“Go”) and indirect (“No-Go”) pathway, CINs stand to allow cognitive flexibility by modulating “Go” and “No-Go” actions (17, 25). Several studies have demonstrated that alcohol preferentially increases glutamatergic transmission in D1-MSNs, but not in D2-MSNs, an effect that potentiates the “Go” pathway (23, 24, 27–29). However, it remains unclear how alcohol affects CIN-mediated modulation of D1- and D2-MSNs.

In the present study, we demonstrated that chronic alcohol intake and withdrawal impaired cognitive flexibility in reversing action-outcome contingency. We found that chronic alcohol intake reduced thalamic inputs to CINs. In the meantime, chronic alcohol consumption led to reduced pause responses of CINs along with increased spontaneous firing activities. Moreover, chronic alcohol intake impaired both CIN-mediated inhibition of glutamatergic transmission in D1-MSNs and CIN-mediated short-term facilitation of glutamatergic transmission in D2-MSNs. These results indicate that alcohol consumption is associated with distinctive CIN-mediated changes in different MSN circuits, providing a potential neural mechanism driving the inflexible drinking underlying alcohol use disorder.

## Results

### Chronic alcohol consumption and withdrawal impair reversal of operant learning in rats

Thalamic inputs to DMS CINs have been implicated in the reversal of instrumental learning (9, 10, 30). We thus examined whether chronic alcohol intake and withdrawal affected the acquisition and reversal of action-outcome contingencies. Rats that had been exposed to water (controls) or 20% alcohol using an intermittent-access 2-bottle choice drinking procedure (24, 31–34) for 8 weeks were trained to learn two action-outcome contingencies involving food pellets or sucrose solution (9, 35) (Fig. 1A). The water and alcohol groups both acquired action-outcome contingencies during the increased-effort training schedule (Fig. 1B). The total number of lever presses was slightly lower in the alcohol group than in the water group, but this difference was not statistically significant (Fig. 1B; *F*_(1, 22)_ = 3.55, *p* = 0.07). Cumulative lever presses during the last session of the initial learning period did not differ between the two groups (Fig. 1C; *F*_(1, 22)_ = 0.13, *p* > 0.05).

**Figure 1.**
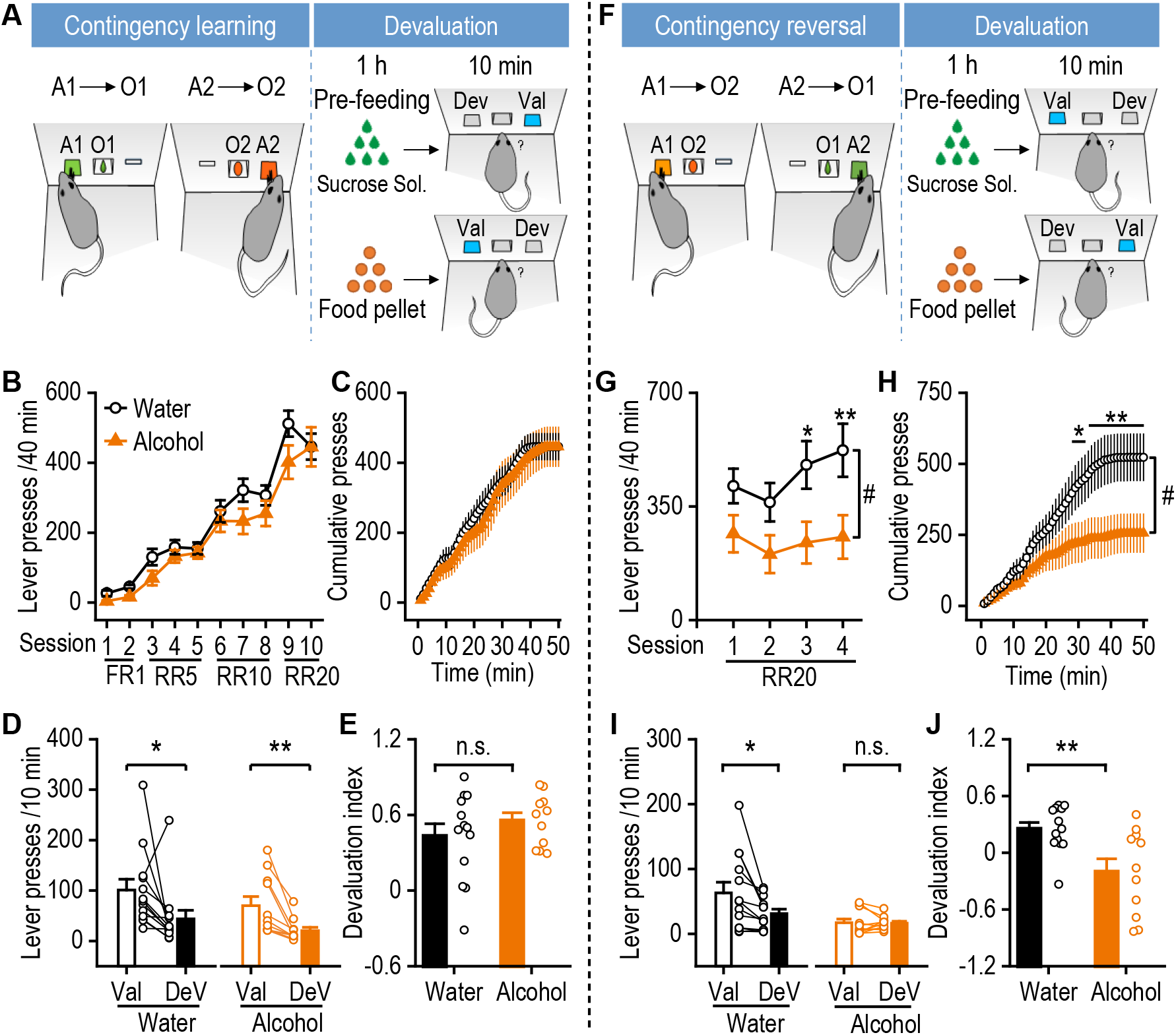
Chronic alcohol intake impairs reversal of instrumental learning. **(A)** Schematic diagram depicting the instrumental learning procedure and subsequent devaluation testing. Long-Evans rats were trained to consumed 20% alcohol using the intermittent-access 2-bottle choice drinking procedure or received water for 8 weeks prior to operant training. Left, alcohol-exposed rats and water controls were trained to acquire the first set of action-outcome (A-O) contingencies, where pressing one of two available levers (A1 or A2) provided a reward of either a food pellet (O1) or sucrose solution (O_2_). Right, outcome-specific devaluation tests involved pre-feeding the rats with O1 or O_2_ for 1 h prior to extinction choice testing (A1 versus A2). The lever associated with the pre-fed reward was defined as devalued (DeV), and the other lever was defined as valued (Val). **(B)** The alcohol and water groups showed no significant difference in total lever presses during the acquisition of the initial contingencies, moving from a fixed ratio 1 (FR1) schedule to random ratio 5 (RR5), RR10, and RR20 schedules as indicated; two-way RM ANOVA, n = 13 rats (Water) and 11 rats (Alcohol). **(C)** The alcohol and water groups showed no difference in cumulative lever presses during the last initial training session (session 10); two-way RM ANOVA, n = 13 rats (Water) and 11 rats (Alcohol). **(D)** Outcome-specific devaluation testing showed that both water and alcohol groups pressed the DeV lever significantly fewer times than the Val lever; \**p* < 0.05 (Water) and \*\**p* < 0.01 (Alcohol) by paired *t* test, n = 13 rats (Water) and 11 rats (Alcohol). **(E)** The devaluation index, defined as (Val - DeV)/(Val + DeV), did not differ significantly between the two groups; n.s., not significant by unpaired *t* test, n = 13 rats (Water) and 11 rats (Alcohol). **(F)** Schematic diagram showing the next round of instrumental learning, with reversed contingencies and subsequent devaluation testing. Left, alcohol rats and water controls were trained to acquire the reversed set of A-O contingencies for 4 days using the RR20 schedule. Right outcome-specific devaluation testing was performed as described above. **(G)** The alcohol group showed significantly reduced lever pressing during the reversed contingency training sessions, as compared to the water group; #*p* < 0.05 by two-way RM ANOVA; \**p* < 0.05, \*\**p* < 0.01 versus the same session in the alcohol group by *Tukey post-hoc* test; n = 13 rats (Water) and 11 rats (Alcohol). **(H)** The alcohol group showed significantly fewer cumulative lever presses in the last reversal learning session (session 4), as compared to the water group; #*p* < 0.05 by two-way RM ANOVA; \**p* < 0.05, \*\**p* < 0.01 for group comparisons at the indicated time points by *Tukey post-hoc* test; n = 12 rats (Water) and 10 rats (Alcohol). **(I)** Outcome-specific devaluation after the reversed A-O contingency learning showed that the water group interacted less with the DeV lever, but this devaluation was not observed in the alcohol group; \**p* < 0.05 (Water) and n.s., not significant, *p* > 0.05 (Alcohol) by paired *t* test, n = 13 rats (Water) and 11 rats (Alcohol). **(J)** The devaluation index was significantly lower in the alcohol group than in the water group; \*\**p* < 0.01 by unpaired *t* test; n = 13 rats (Water) and 11 rats (Alcohol).

After the initial acquisition of this task, we investigated the sensitivity to outcome devaluation. To achieve this goal, animals were fed with either food pellets or sucrose solution before receiving extinction training, where lever presses were monitored. We found that both alcohol-drinking and water control rats significantly decreased their presses on the outcome-satiated (devalued) lever (Fig. 1D; *t*_(12)_ = 2.20, *p* < 0.05 for water group; *t*_(10)_ = 3.71, *p* < 0.01 for alcohol group). Analysis of the devaluation index (the difference between the proportion of non-devalued and devalued lever presses) did not identify any statistically significant difference between the degree of goal-directed versus habitual behavior in the alcohol-drinking and water control rats (Fig. 1E; *t*_(22)_ = −1.02, *p* > 0.05). These results indicated that alcohol-drinking and water control rats showed similar levels of goal-directed behavior.

Next, we examined the flexibility of the rats’ responses to a change in the action-outcome contingency. We reversed the relationship between action and outcome so that pressing the lever previously used to access sucrose solution now led to the delivery of food pellets and vice versa (Fig. 1F). Following this contingency reversal, the total lever presses were significantly lower in the alcohol group than in the control group (Fig. 1G; *F*_(1,22)_ = 6.28, *p* < 0.05). Cumulative lever presses were also lower in the alcohol-drinking rats than in water controls during the last session of reversal training (Fig. 1H; *F*_(1,20)_ = 4.68, *p* < 0.05). These results indicated that chronic alcohol intake and withdrawal (at least 10 d) impaired reversal learning in this task.

Lastly, our analysis of the relative contributions of goal-directed versus habitual behavior following contingency reversal showed that the alcohol group pressed indiscriminately on devalued and non-devalued levers, whereas the water control rats still favored the non-devalued lever (Fig. 1I; *t*_(12)_ = 2.87, *p* < 0.05 for water group; *t*_(10)_ = 0.18, *p* > 0.05 for alcohol group). The devaluation index was, therefore, significantly lower in alcohol-drinking rats, as compared to their water controls (Fig. 1J; *t*_(22)_ = 3.14, *p* < 0.01). We also compared the difference between the first and second devaluation indices in the two study groups; the alcohol group showed a significantly larger decrease than did the water group (Supplementary Fig. 1; *t*_(22)_ = 2.88, *p* < 0.01). These results indicated that the water controls maintained a goal-directed strategy in response to the new action-outcome association. However, alcohol-drinking rats failed to do so and instead used a strategy more consistent with habitual behavior, suggesting that chronic alcohol intake and withdrawal impaired cognitive flexibility in response to changes in action-outcome associations in rats.

### Chronic alcohol consumption reduces glutamatergic thalamostriatal inputs onto DMS CINs

The striatum receives major glutamatergic inputs from both the cortex and thalamus. Reduced flexibility in reversal learning is known to be associated with thalamostriatal transmission in DMS CINs (9, 17, 36). We next investigated whether alcohol consumption altered thalamic inputs to DMS CINs. To selectively induce thalamostriatal transmission, we expressed channelrhodopsin 2 (ChR2) in thalamic inputs (Fig. 2A) by crossing transgenic mice expressing Cre recombinase under the control of the vesicular glutamate transporter 2 (VGluT2) promoter (VGluT2-Cre mice) with transgenic mice with Cre-dependent ChR2-eYFP expression (Ai32 mice) (37). This cross produced VGluT2-Cre;Ai32 mice. Previous studies in VGluT2-Cre mice reported that VGluT2-expressing inputs to the striatum mainly arose from the thalamus (38, 39).

**Figure 2.**
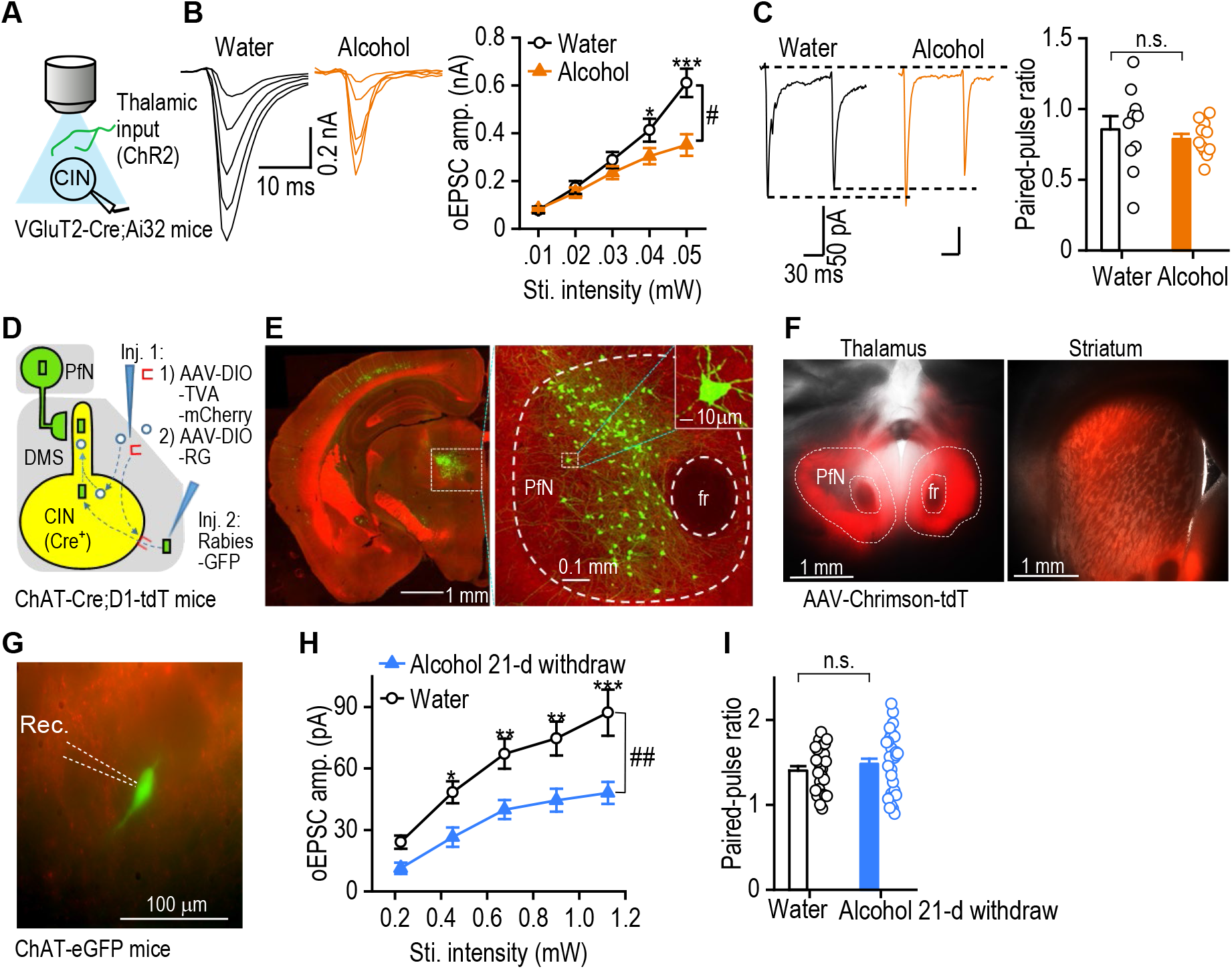
Chronic alcohol consumption reduces thalamostriatal glutamatergic inputs onto DMS CINs. **(A)** Schematic diagram showing light stimulation of ChR2-expressing thalamic inputs and whole-cell recording of CINs. **(B)** Chronic alcohol consumption suppressed thalamostriatal transmission onto CINs in DMS slices prepared 24 h after the last alcohol exposure. Left and middle, sample traces of EPSCs evoked by the indicated optical stimulation (oEPSCs). Right, input-output curves of oEPSC amplitudes in CINs from mice exposed to alcohol or water; #*p* < 0.05 by two-way RM ANOVA; \**p* < 0.05, \*\*\**p* < 0.001 versus the same stimulation intensity in the alcohol group by *Tukey post-hoc* test, *n* = 11, 4 (Water) and 13, 3 (Alcohol). **(C)** Chronic alcohol consumption did not alter the glutamate release probability at thalamostriatal synapses. Left and middle, representative traces of oEPSCs induced by paired-pulse optical stimulations in the alcohol and water groups. Right, the paired-pulse ratios in the indicated groups; *p* > 0.05 by unpaired *t* test, *n* = 10, 3 (Water) and 11, 3 (Alcohol). **(D)** Schematic showing viral infusions. We injected rabies helper viruses (AAV-DIO-TVA-mCherry and AAV-DIO-RG) into the DMS of ChAT-Cre;D1-tdT mice, leading to selective expression of TVA and RG in CINs (Inj. 1). D1-tdT was used to show the background. After 2 weeks, we injected rabies-GFP into the same DMS site (Inj. 2). This approach caused selective TVA-mediated infection of CINs with rabies-mCherry, followed by RG-mediated retrograde transsynaptic infection of presynaptic neurons, including those in the thalamic parafascicular nucleus (PfN). **(E)** Sample coronal images showing that many rabies-GFP-labeled PfN neurons projected to DMS CINs. Similar results were observed in 4 mice; fr, fasciculus retroflexus. **(F)** Optical images of tdT fluorescence in striatal slices from ChAT-eGFP mice that were infused with AAV-Chrimson-tdT in the PfN of the thalamus. **(G)** High magnification optical image showing an GFP expressing CIN and its surround tdT fluorescence. **(H)** Input-output curves of oEPSC amplitudes in CINs from mice injected with AAV-Chrimson-tdT and exposed to alcohol or water; ##*p* < 0.01 by two-way RM ANOVA; \**p* < 0.05, \*\**p* < 0.01, \*\*\**p* < 0.001 versus the same stimulation intensity in the alcohol group by *Tukey post-hoc* test, *n* = 15, 4 (Water) and 14, 4 (Alcohol). **(I)** Paired-pulse ratios in mice injected with AAV-Chrimson-tdT and exposed to alcohol or water; *p* > 0.05 by unpaired *t* test, *n* = 27, 5 (Water) and 32, 5 (Alcohol).

CINs are easily distinguished from other striatal cell types because they have a large diameter and unique electrophysiological characteristics (40, 41). We thus distinguished CINs from MSNs by their larger size, spontaneous firing (Supplementary Fig. 2A), higher resting membrane potential, characteristic voltage sag in response to hyperpolarizing current injection, and greater excitability in response to depolarizing current injection (Supplementary Fig. 2B, resting membrane potentials: *t*_(10)_ = 4.75, *p* < 0.001). Interestingly, repetitive light-mediated stimulation of thalamic inputs in VGluT2-Cre;Ai32 mouse slices evoked distinct patterns of excitatory postsynaptic potentials (EPSPs) in CINs and MSNs. We found that the second EPSP was larger than the first EPSP in CINs, while MSNs showed the opposite pattern (Supplementary Fig. 2C; *t*_(10)_ = 6.87, *p* < 0.001). We used a combination of these approaches to identify CINs when these neurons did not express fluorescent proteins.

We then explored how chronic alcohol intake influenced thalamostriatal glutamatergic transmission onto DMS CINs. VGluT2-Cre;Ai32 mice were trained to consume 20% alcohol for 8 weeks using the intermittent-access 2-bottle choice drinking procedure (23, 34). Twenty-four hours after the last alcohol exposure, we prepared striatal slices and measured optically-evoked excitatory postsynaptic currents (oEPSCs) in CINs. We found that the oEPSC amplitude was significantly lower in CINs from the alcohol group than those from the water control group (Fig. 2B; *F*_(1, 22)_ = 5.39, *p* < 0.05). This result suggests that chronic alcohol intake reduced thalamostriatal inputs onto DMS CINs. To further investigate the mechanism underlying this reduction, we measured the paired-pulse ratio (PPR) of oEPSCs that were activated by two stimuli, delivered 100 ms apart. This analysis found no difference between the alcohol group and the water group (Fig. 2C; *t*_(19)_ = 0.72, *p* > 0.05). These results suggested that the reduced thalamostriatal transmission to CINs in mice with chronic alcohol exposure was unlikely to be caused by a reduced probability of presynaptic glutamate release.

To further confirm the alcohol-associated suppression of thalamostriatal transmission, we infused an adeno-associated virus (AAV)-Chrimson-tdTomato (tdT) into a thalamic nucleus that is known to project to DMS CINs. Previous studies identified dense inputs to the striatum from multiple thalamic nuclei, including the parafascicular nucleus (PfN) (42, 43). To investigate this, we infused rabies helper viruses into the DMS of ChAT-Cre mice, waited three weeks, and then infused rabies-GFP at the same location 3 weeks later (Fig. 2D). Two Cre-dependent (Flex) AAV serotype 8 vectors were employed as helper viruses; one expressed rabies glycoprotein (RG) (AAV8-DIO-RG), and the other expressed an avian membrane EnvA receptor protein (TVA) and mCherry (AAV8-DIO-TVA-mCherry). This approach produced extensive GFP expression in the PfN (Fig. 2E), indicating dense innervation of DMS CINs by thalamic PfN neurons. Next, we infused AAV-Chrimson-tdT into the PfN of ChAT-eGFP mice and detected the tdT fluorescent signal in the striatum (Fig. 2F). Animals were trained to consume alcohol as described above. Twenty-four hours after the last alcohol exposure, striatal slices were prepared to measure optically evoked excitatory postsynaptic currents (oEPSCs) in CINs. Similar changes in the oEPSCs were observed (Supplementary Fig. 3C; *F*_(1,22)_ = 4.74, *p* < 0.05) as in Figure 2B. We did not observe significant changes in PPR measurements (Supplementary Fig. 3D; *t*_(39)_ = −1.44, *p* > 0.05). Because we had previously observed behavioral deficits weeks after stopping alcohol consumption (Fig. 1), we also measured oEPSCs 21 d after the last alcohol exposure. Similar results were observed at this time-point (oEPSCs: Fig. 2H; *F*_(1,27)_ = 10.91, *p* < 0.01. PPR: Fig. 2I; *t*_(53)_ = −0.72, *p* > 0.05).

Taken together, these data suggest that chronic alcohol consumption causes a long-lasting decrease in thalamostriatal inputs onto DMS CINs.

### Chronic alcohol consumption significantly increases the spontaneous firing of DMS CINs and shortens their pause responses

Having shown that chronic alcohol intake reduced thalamic inputs onto DMS CINs, we asked whether alcohol also altered the spontaneous spiking of these tonically active neurons. We trained ChAT-eGFP mice to consume alcohol for 8 weeks using the intermittent-access 2-bottle choice drinking procedure. CINs were identified by their green fluorescence (Fig. 3A), and spontaneous firing of DMS CINs was measured using cell-attached recording, 24 h and 21 d after the last alcohol exposure. We found that chronic alcohol consumption decreased the inter-spike interval (Fig. 3C) and significantly increased the firing frequency over time (Fig. 3D; *F*_(2, 112)_ = 5.69,, *p* < 0.01). In contrast, our measurement of intrinsic excitability using whole-cell recording did not find any difference in the evoked firing of DMS CINs from the water and alcohol groups (Fig. 3E; *F*_(1, 27)_ = 0.93, *p* > 0.05). These results suggested that chronic alcohol consumption increased the spontaneous activity of DMS CINs.

**Figure 3.**
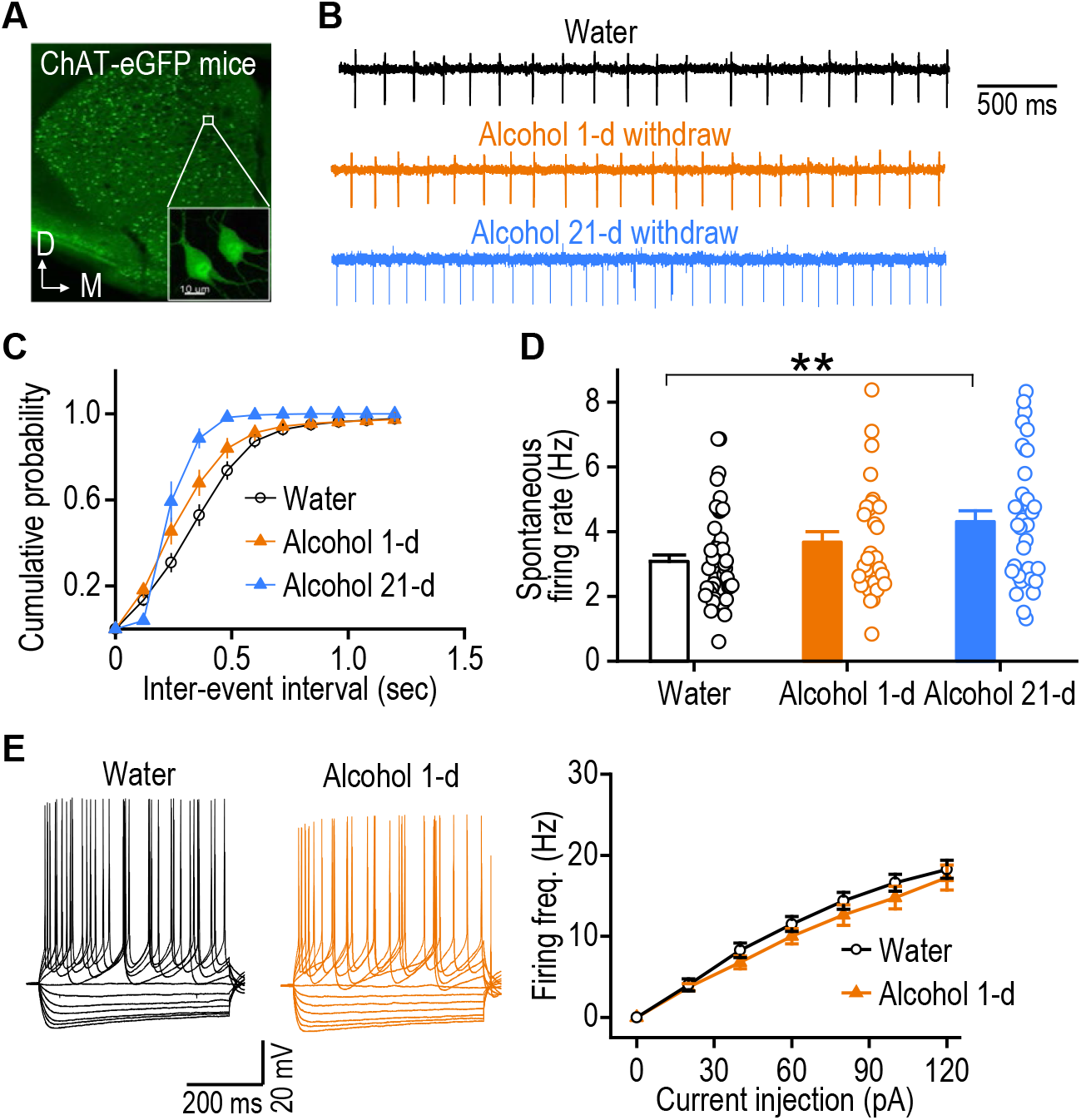
Chronic alcohol consumption increases spontaneous, but not evoked, firing of DMS CINs. ChAT-eGFP mice were trained to consume 20% alcohol for 8 weeks and DMS slices were prepared 24 h and 21-d after the last alcohol exposure. **(A)** Sample image showing green CINs in the striatum. D, dorsal; M, medial. **(B)** Sample traces of spontaneous CIN firing in the water and alcohol groups using the cell-attached recording. **(C)** Cumulative plots of the inter-event intervals and **(D)** the spontaneous firing rates of CINs in the indicated groups; ##*p* < 0.01 by one-way ANOVA, \*\**p* < 0.01 versus water group by *Tukey post-hoc* test; *n* = 49, 7 (Water), 31, 6 (Alcohol 1-d), and 36, 4 (Alcohol 21-d). **(E)** Chronic alcohol did not change evoked CIN firing. Left and middle, sample traces of membrane potentials generated in the indicated groups in response to a series of 500-ms current injections. Right, the input-output relationship between the injected current magnitude and the CIN firing frequency in water and alcohol groups; *p* > 0.05 by two-way RM ANOVA, *n* = 16, 4 (Water) and 13, 3 (Alcohol).

CINs exhibit characteristic burst-pause firing, which is important for regulating MSN activity. Next, we investigated the effects of chronic alcohol intake on the burst-pause firing of CINs. To induce burst-pause response of CINs, we expressed ChR2 in CINs by crossing transgenic mice expressing Cre recombinase under the control of choline acetyltransferase (ChAT) promoter (ChAT-Cre mice) with transgenic mice with Cre-dependent ChR2-eYFP expression (Ai32 mice)(37). ChAT-Cre;Ai32 mice were trained to consume 20% alcohol for 8 weeks using the intermittent-access 2-bottle choice drinking procedure. Twenty-four hours after the last alcohol exposure, we prepared striatal slices and measured optically evoked burst-pause responses of DMS CINs. We found that the pause duration was significantly shorter in CINs from the alcohol group than those from the water control group using cell-attached recording (Fig. 4 A, B; *t*_(40)_ = 2.32, *p* < 0.05). We also observed similar results with whole-cell recording (Fig. 4 C, D; *t*_(31)_ = 2.06, *p* < 0.05).

**Figure 4.**
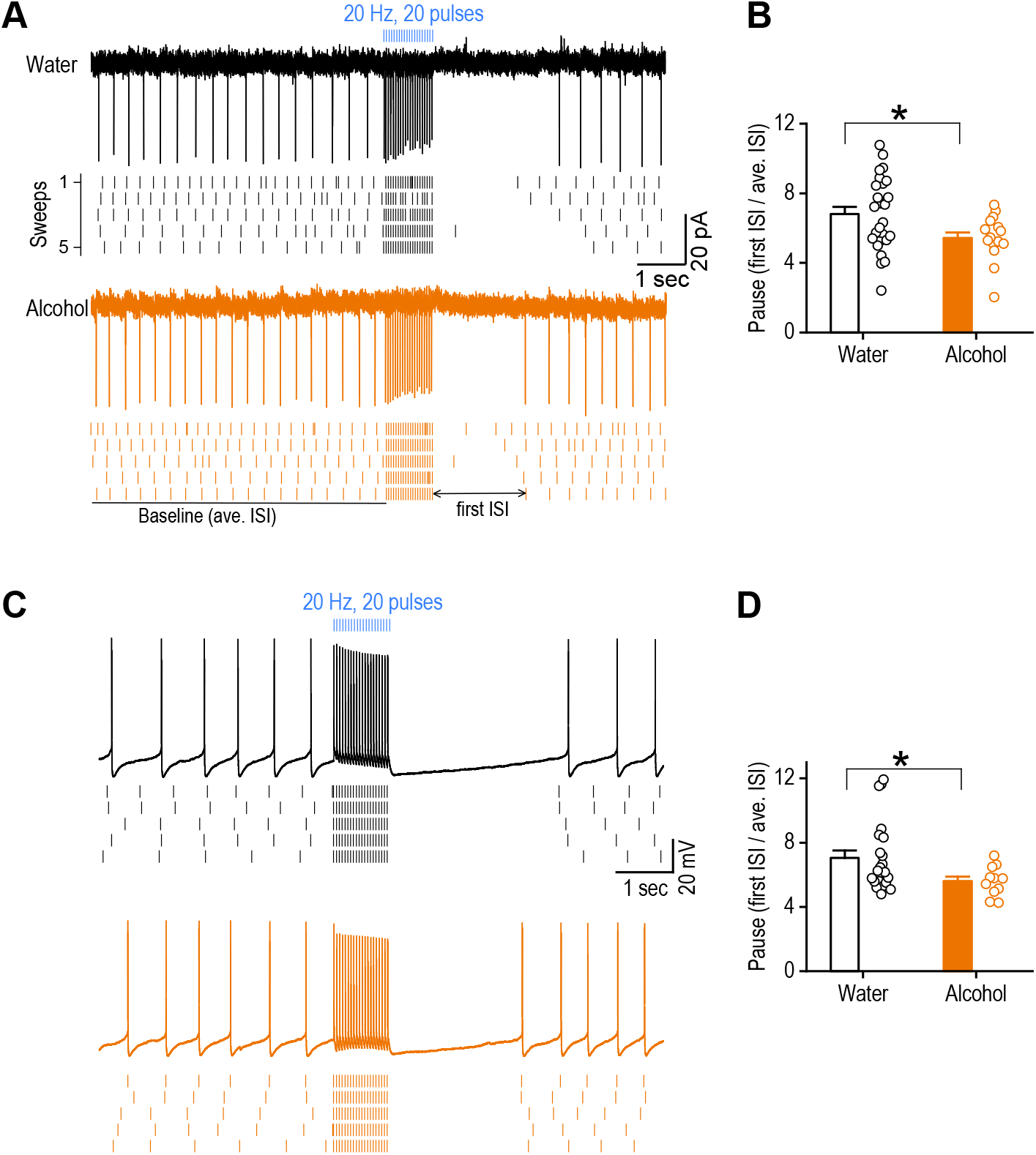
Chronic alcohol consumption shortens pause response of DMS CINs. ChAT-Cre;Ai32 mice were trained to consume 20% alcohol for at least 8 weeks. Then DMS slices were prepared 24 h after last alcohol exposure and optically-evoked burst-pause responses of CINs were measured. **(A)** Sample traces of burst-pause responses of a CIN from the water (top) and alcohol (bottom) groups using the cell-attached recording. ISI: inter-spike interval. **(B)** The pause durations in the indicated groups; *p* < 0.05 by unpaired *t* test, *n* = 26, 5 (Water) and 16, 4 (Alcohol). The pause duration is defined by the first ISI right after optical stimulation divided by baseline average ISI before the optical stimulation. **(C)** Sample traces of burst-pause responses of a CIN from the water (top) and alcohol (bottom) groups using whole-cell recording. **(D)** The pause durations in the indicated groups; *p* < 0.05 by unpaired *t* test, *n* = 22, 5 (Water) and 11, 3 (Alcohol).

### Chronic alcohol consumption impairs CIN-induced suppression of NMDAR-mediated glutamatergic transmission in DMS D1-MSNs

CINs regulate flexible behaviors by modulating MSN activity. After characterizing the effects of chronic alcohol consumption on DMS CIN activity, we investigated how alcohol intake might affect CIN-mediated modulation of MSNs, leading to changes in striatal output. In striatal circuits, muscarine is known to modulate NMDA receptor (NMDAR)-mediated synaptic responses in D1-MSNs by acting on muscarinic M4 receptors (26). We, therefore, examined whether endogenous acetylcholine release induced by optogenetic excitation of CINs altered NMDAR-EPSCs in DMS D1-MSNs. To achieve this, we generated triple transgenic ChAT-Cre;Ai32;D1-tdT mice, in which CINs expressed ChR2-eYFP and D1-MSNs contained tdT (Fig. 5A). Stimulating electrodes were placed within the striatum to elicit glutamatergic transmission, and we patched the (red) D1-MSNs and excited CINs by delivering blue light through the objective lens (Fig. 5B). After NMDAR-mediated EPSCs were recorded for 5 min (baseline), blue light (2 ms, 10 pulses at 15 Hz) was delivered 1 sec prior to each electrical stimulation, and EPSCs were continuously monitored for 10 min (Fig. 5C). We found that optogenetic excitation of CINs significantly reduced the NMDAR-EPSC amplitude in D1-MSNs (Fig. 5D; *t*_(6)_ = 6.13, *p* < 0.001). We further confirmed that this effect was mediated by muscarinic M4 receptors, as subsequent application of an antagonist of this receptor, PD 102807 (1 μM) (44), completely abolished the CIN-mediated suppression of NMDAR-EPSCs (Fig. 5D; *t*_(6)_ = −4.64, *p* < 0.01). We found that chronic alcohol consumption completely abolished this CIN excitation-induced suppression of NMDAR-EPSCs in D1-MSNs (Fig. 5E; *t*_(6)_ = −0.68, *p* > 0.05). Taken together, these data indicated that excitation of DMS CINs activated muscarinic M4 receptors to suppress NMDAR-EPSCs in DMS D1-MSNs and that chronic alcohol consumption attenuated this suppression.

**Figure 5.**
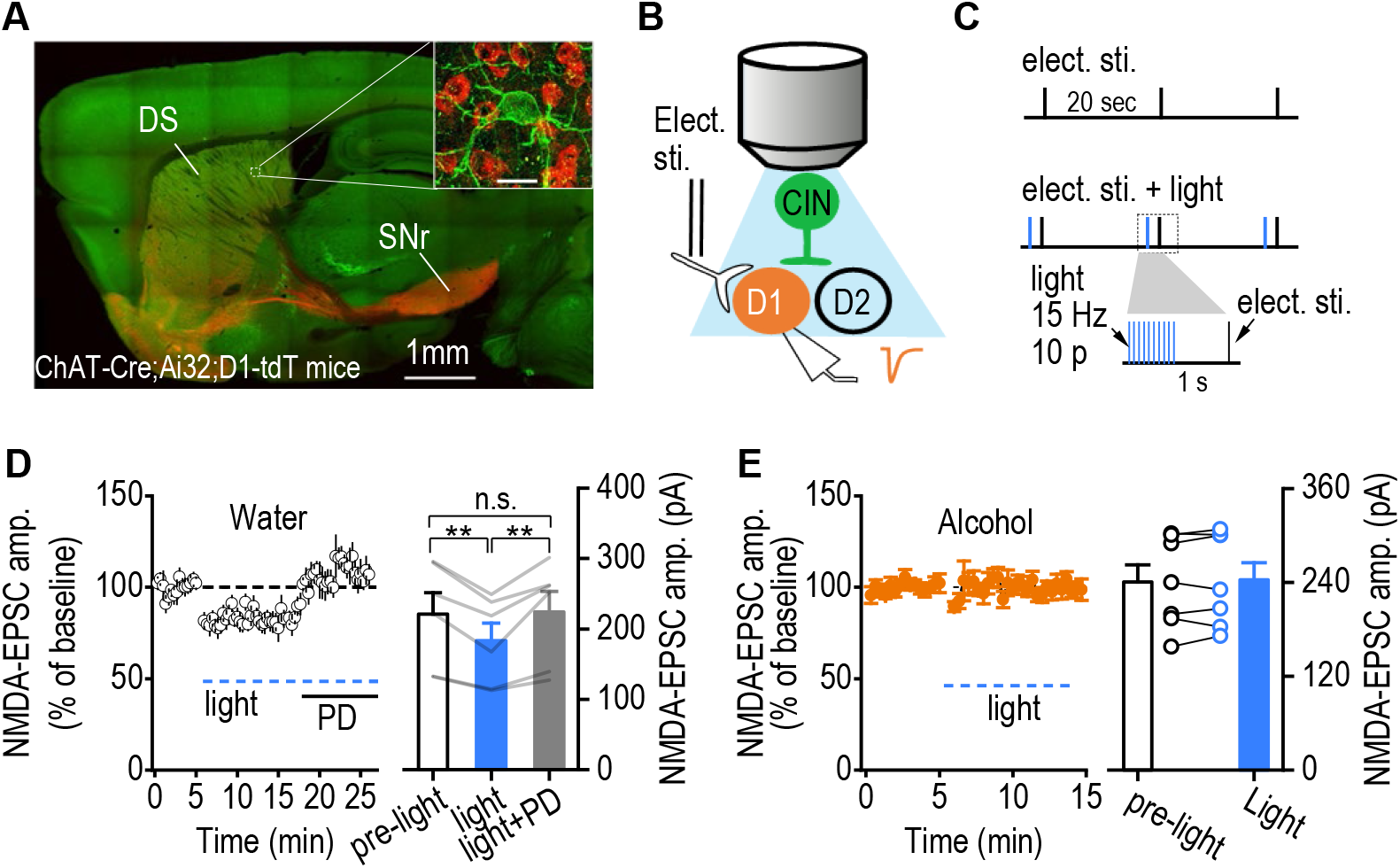
Chronic alcohol consumption impairs CIN-mediated suppression of glutamatergic transmission in DMS D1-MSNs. **(A)** Sample image of a sagittal section from a ChAT-Cre;Ai32;D1-tdT mouse. Inset shows a green CIN with several red D1-MSNs (scale bar: 20 μm). **(B)** Schematic illustration of the electrical and optical stimulation and selective recording of D1-MSNs. The stimulating electrodes were placed in the DMS close to the recording pipette. **(C)** Schematic of the electrical and light stimulation protocols. Electrical stimulation (top) was delivered every 20 sec, 1 sec after the delivery of a burst of 473-nm light (2 ms of 10 pulses at 15 Hz) (middle and bottom). **(D)** The amplitude of NMDAR-mediated EPSCs before light stimulation, during light stimulation, and during light stimulation in the presence of the muscarinic M4 antagonist, PD 102807 (PD, 1 μM), showed that optogenetic excitation of DMS CINs caused an M4 receptor-dependent suppression of NMDAR activity in D1-MSNs; \*\**p* < 0.01, unpaired *t* test, *n* = 7, 5 per group. **(E)** Chronic alcohol consumption abolished CIN-induced suppression of NMDAR-EPSCs; *p* > 0.05 by paired *t* test, *n* = 7, 4 per group.

### Chronic alcohol consumption compromises CIN-mediated short-term facilitation of glutamatergic transmission in DMS D2-MSNs

Having found that chronic alcohol consumption impaired CIN-mediated regulation of glutamatergic transmission in D1-MSNs, we next examined whether it altered CIN-mediated regulation of glutamatergic transmission in another major MSN type, the D2-MSN. We employed ChAT-Cre;Ai32;D1-tdT mice, in which putative D2-MSNs were identified as non-fluorescent (Fig. 6A). Thalamic stimulation of cholinergic activity has been shown to cause short-term facilitation of AMPAR-EPSPs in D2-MSNs (17). We thus recorded electrically-evoked AMPAR-EPSPs in D2-MSNs using the current-clamp recording. Five EPSPs were measured before and 1 sec after light-mediated stimulation of CINs in mice that had been exposed to alcohol or water only. Compared to amplitudes recorded before light stimulation, we found that direct light stimulation (15 Hz, 10 pulses, 1 sec before electrical stimulation) of CINs caused short-term facilitation of EPSP amplitudes in the water group (Fig. 6B; *F*_(1,8)_ = 5.66, *p* < 0.05), as expected. Interestingly, there was also a main effect of pulse number (*F*_(4,32)_ = 3.89, *p* < 0.05), in that later electrical pulses generated higher relative EPSP amplitudes than earlier pulses (Fig. 6B; versus pulse 1: *q* = 4.53, *p* < 0.05 (pulse 2); *q* = 6.67, *p* < 0.001 (pulse 4); *q* = 7.41, *p* < 0.001 (pulse 5)). In contrast, light stimulation of CINs failed to potentiate the EPSP amplitudes in the alcohol group (Fig. 6C; *F*_(1, 11)_ = 0.91, *p* > 0.05). These results demonstrated that chronic alcohol consumption compromised CIN-mediated short-term facilitation of AMPAR-mediated transmission in DMS D2-MSNs.

**Figure 6.**
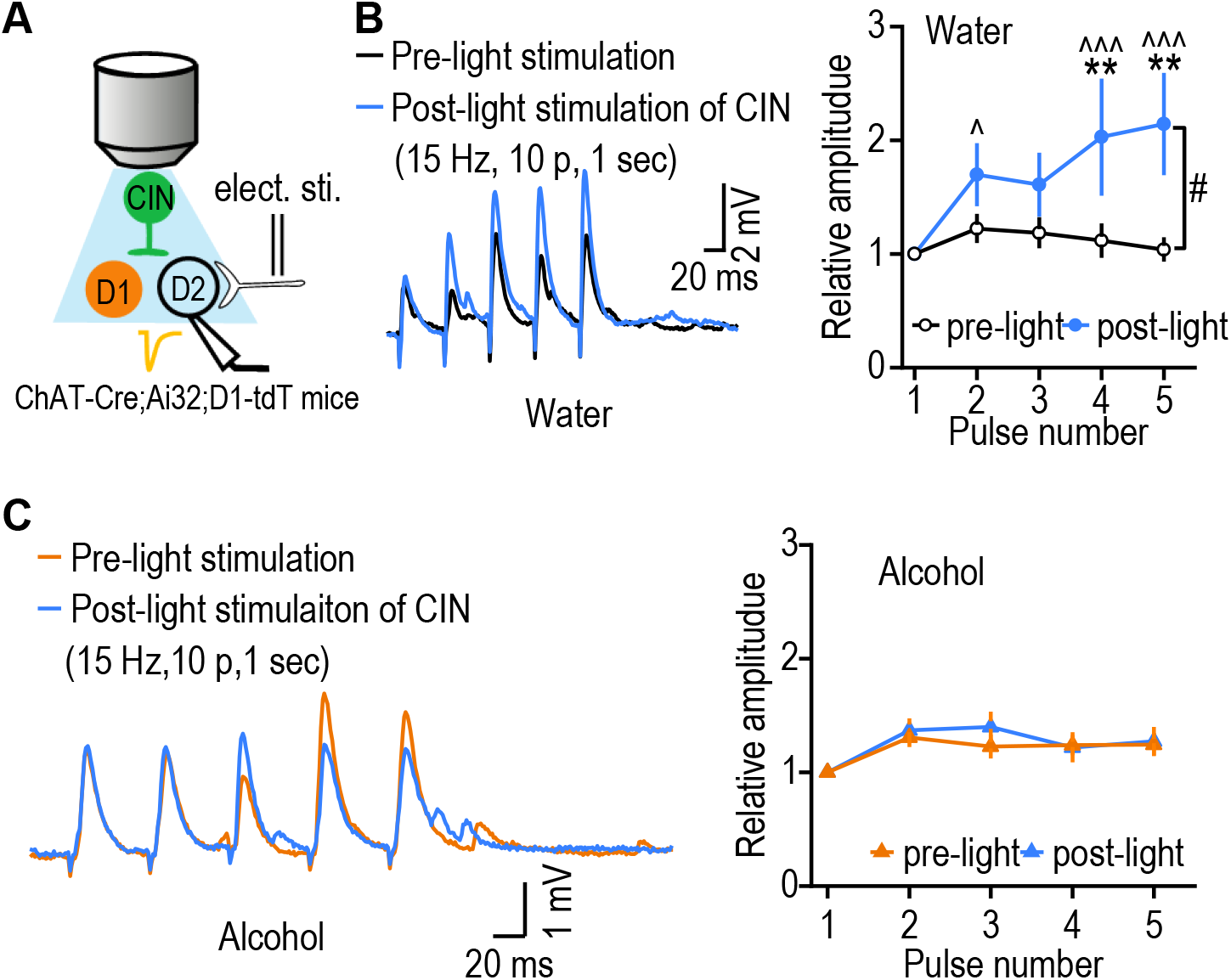
Chronic alcohol intake compromises CIN-mediated short-term facilitation of glutamatergic transmission in DMS D2-MSNs. **(A)** Schematic illustration of the electrical and light stimulation and whole-cell recording of D2-MSNs in ChAT-Cre;Ai32;D1-tdT mice. Putative D2-MSNs were identified by their absence of fluorescence. **(B)** Left, sample traces showing that electrical stimulation led to five EPSPs in D2-MSNs before and after light-mediated excitation of CINs. Electrical stimulation was delivered every 20 sec, 1 sec after the delivery of a bust of 473-nm light (2 ms of 10 pulses at 15 Hz). Right, calculation of the relative amplitudes of five EPSPs detected short-term facilitation in water control mice after light-mediated excitation of CINs. EPSPs were normalized to the first one; #*p* < 0.05 by two-way RM ANOVA; \*\**p* < 0.01 versus the same pulse number in the pre-light group by *Tukey post-hoc* test; ^*p* < 0.05, ^^^*p* < 0.001 versus pulse number 1 within the post-light group by *Tukey post-hoc* test; *n* = 9, 6 per group. **(C)** Left, sample traces showing the EPSPs before and after light stimulation of CINs in the alcohol group. Right, calculation of the relative amplitudes of EPSPs in the alcohol group did not identify any change after light stimulation of CINs; *p* > 0.05; two-way RM ANOVA, *n* = 12, 4 per group.

### The alcohol-induced impairment of reversal learning is rescued by in vivo optogenetic induction of long-term potentiation of PfN-to-CIN transmission

The above evidence points to the key roles of DMS CINs in mediating the detrimental effect of chronic alcohol intake on cognitive flexibility. Lastly, we aimed to alleviate this detrimental effect by manipulating the PfN➔CIN connectivity. It has been shown that a global enhancement of the neuronal activity of CINs through pharmacogenetics failed to rescue the impairment of reversal learning in aged mice (10), indicating the need for a more targeted modulation of CINs by thalamostriatal processes. Therefore, we infused AAV-Chronos-GFP into the PfN and AAV-FLEX-Chrimson-tdT into the DMS for selective manipulation of PfN➔CIN synapses. Optical fibers were implanted into DMS (Fig. 7A). After recovery from surgery, rats were trained using the schedule described in Figure 1. Once the rats acquired the initial A-O contingencies (Fig. 7C, D), they were divided into two groups: Alcohol-Opto group received time-locked light stimulation (Fig. 7B) during the reversal learning; Alcohol-Sham group underwent the same procedure as Alcohol-Opto. group except the light lasers were not turned on. Both groups showed similar acquisition of initial A-O contingencies and initial devaluations (Supplementary Fig. 5). During reversal training, we delivered optogenetic high-frequency stimulation (oHFS) of PfN inputs and optogenetic depolarization (oPSD) of DMS CINs, a dual-channel optogenetic protocol that we recently developed to induce long-term potentiation (LTP) in vivo (24). We found that there was no significant difference in terms of lever presses between the two groups (Fig. 7E; *F*_(1,16)_= 0.002, *p* > 0.05). However, our analysis of the relative contributions of goal-directed versus habitual behavior following contingency reversal showed that the sham group pressed more devalued levers, indicating habitual behavior carrying over from initial learning; whereas the light stimulation group still favored the non-devalued lever, indicating new goal-directed behavior (Fig. 7F; *t*_(8)_ = −1.52, *p* > 0.05 for the sham group; *t*_(9)_ = 1.91, *p* < 0.05 for light stimulation group). The devaluation index was therefore significantly higher in light-stimulated rats, as compared to their sham controls (Fig. 7G; *t*_(17)_ = −2.23, *p* < 0.05). These results indicated that the alcohol-induced impairment of cognitive flexibility was restored by selectively potentiating thalamic inputs onto DMS CINs.

**Figure 7.**
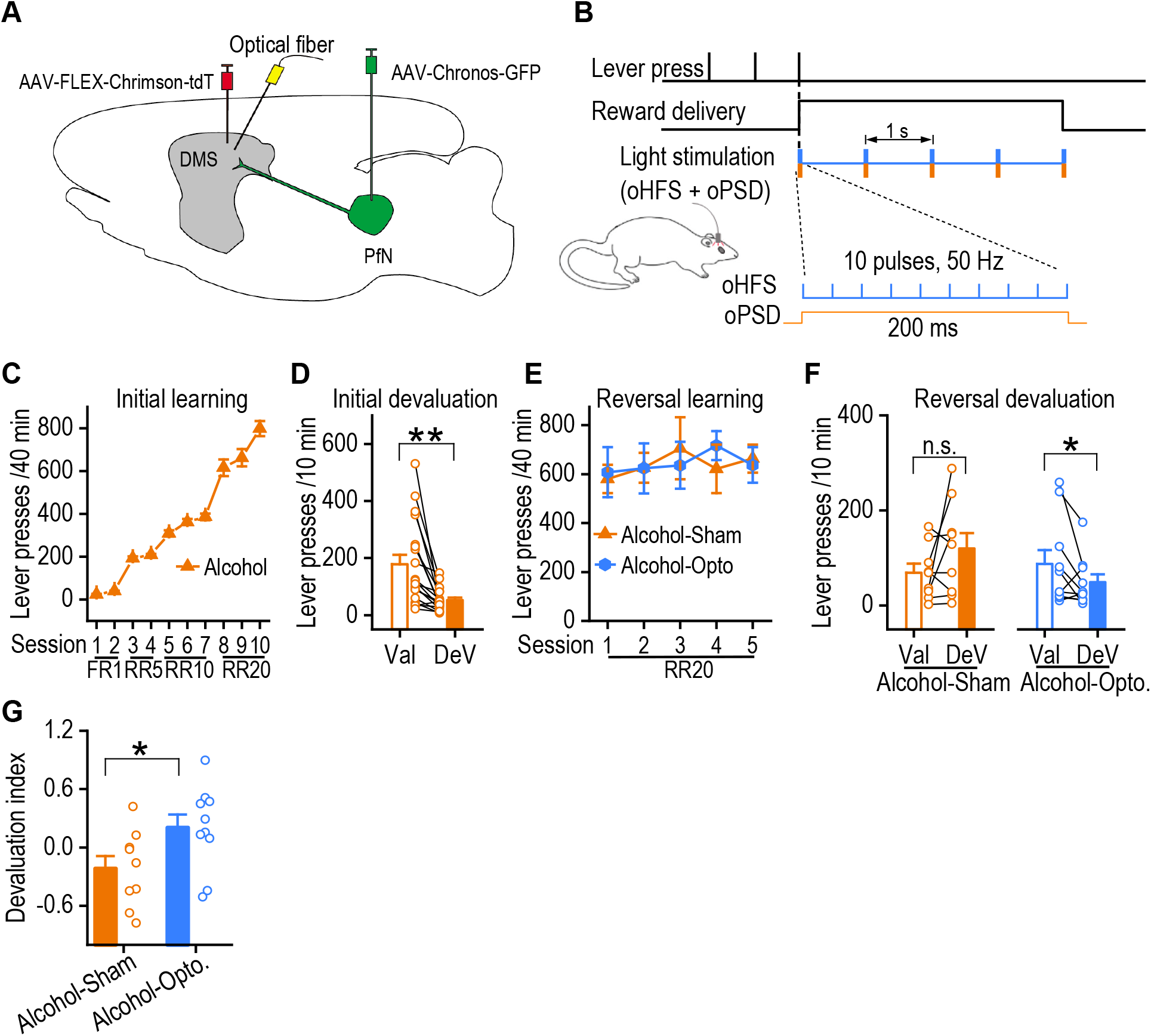
Optogenetic stimulation PfN-to-CIN pathway in the DMS rescues the impairment of reversal learning. **(A)** Schematic diagram of viruses injection and optical fiber implantation. ChAT-Cre rats were bilaterally injected with AAV-FLEX-Chrimson-tdT and AAV-Chronos-GFP into DMS and PfN, respectively. Optical fibers were bilaterally implanted into DMS. After recovery from surgery rats were trained by the schedule as Figure 1. **(B)** Optical light stimulation protocol used during the reversal learning. Briefly, rats press the lever to get the reward. Light stimulation is time-locked to reward delivery. Light stimulation contains five repeats of dual light stimulus within 5 s reward delivery period. Each repeat consists of blue light high frequency stimulation (10 pulses, 50 Hz) and yellow light (continuous, 200 ms) delivering at the same time. oHFS: optical high-frequency stimulation; oPSD: optical postsynaptic depolarization. **(C)** The initial acquisition learning curve. n = 19. **(D)** Outcome-specific devaluation testing showed that rats pressed the DeV lever significantly fewer times than the Val lever; \*\**p* < 0.01 by paired *t* test, n = 19. **(E)** There was no significantly difference in terms of lever pressing between two groups during the reversed contingency training sessions; #*p >* 0.05 by two-way RM ANOVA; n = 9 rats (Alcohol-Sham) and 10 rats (Alcohol-Opto). **(F)** Outcome-specific devaluation after reversed A-O contingency learning showed that the sham group still interacted more with the DeV lever (which is Val lever during initial learning), while the group received light stimulation showed successful devaluation after the reversed A-O contingency; n.s., not significant, *p* > 0.05 and \**p* < 0.05 by paired *t* test, n = 9 rats (Alcohol-Sham) and 10 rats (Alcohol-Opto). **(G)** The devaluation index was significantly higher in the opto group than in the sham group; \**p* < 0.05 by unpaired *t* test; n = 9 rats (Alcohol-Sham) and 10 rats (Alcohol-Opto).

## Discussion

In this study, we demonstrated that chronic alcohol exposure and withdrawal reduced goal-directed cognitive flexibility and caused a long-lasting suppression of thalamostriatal inputs onto DMS CINs and a shortened pause response along with the increased spontaneous firing of these neurons. Furthermore, chronic alcohol consumption and withdrawal impaired CIN-mediated downregulation of glutamatergic transmission in D1-MSNs, as well as CIN-mediated short-term upregulation of glutamatergic transmission in D2-MSNs. Our data suggest that chronic alcohol consumption compromises the thalamostriatal regulation of glutamatergic transmission in MSNs via CINs (Fig. 8), providing insight into how chronic alcohol consumption changes from casual, flexible drinking to compulsive intake.

**Figure 8.**
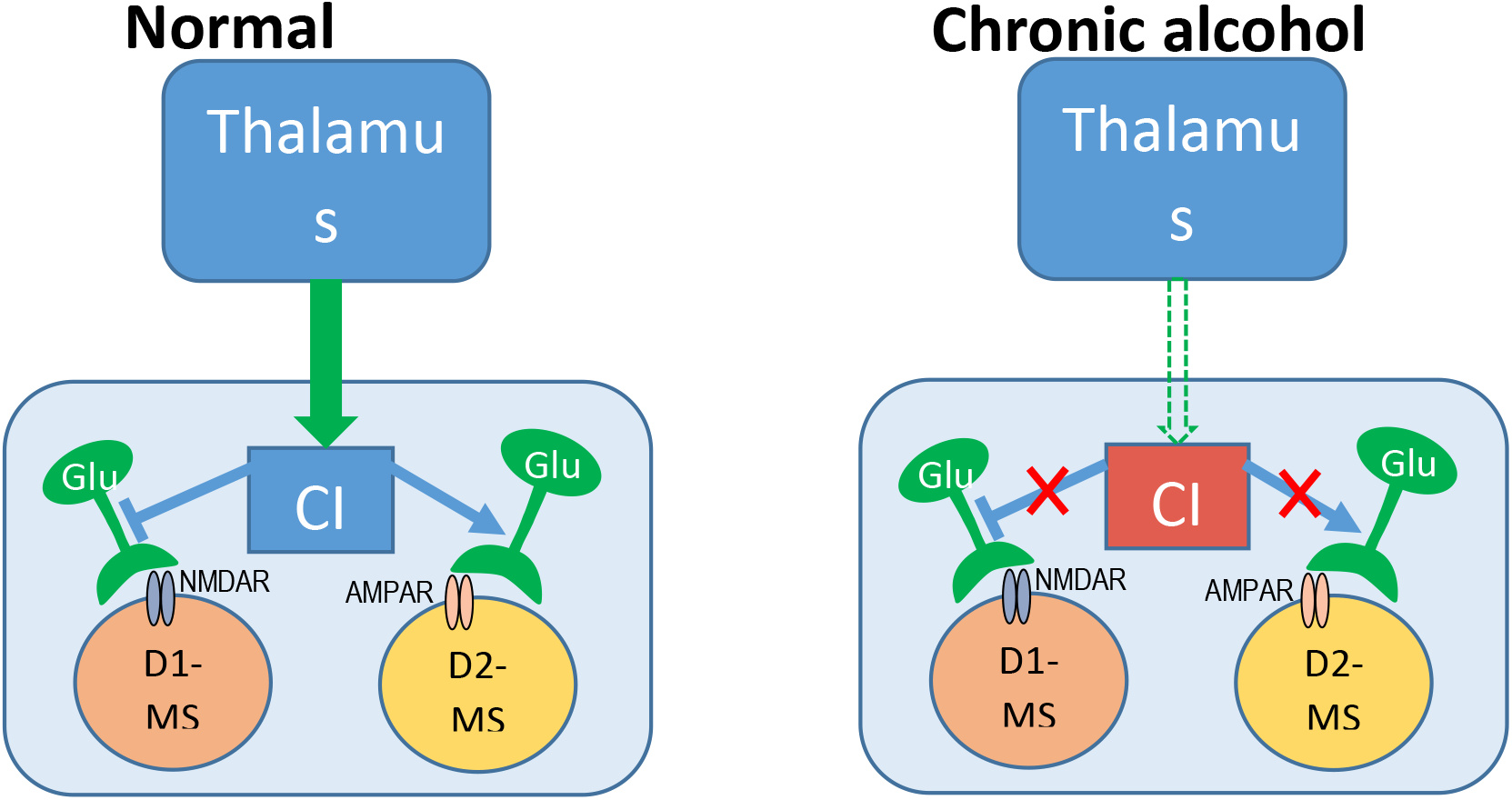
Schematic diagram showing the effects of chronic alcohol intake on the thalamic inputs to CINs and their modulation of glutamatergic transmission to D1-MSN and D2-MSN in the striatum. Chronic alcohol consumption reduces thalamic excitatory inputs to DMS CINs and increases their spontaneous firing, which makes them less prone to be modulated by external signals. In the meantime, the CIN-mediated inhibition of glutamatergic transmission in D1-MSNs and the CIN-mediated short-term facilitation of glutamatergic transmission in D2-MSNs are comprised after chronic alcohol intake, which could change striatal outputs and lead to behavioral inflexibility.

In individuals with alcohol use disorder, a progressive loss of cognitive flexibility eventually results in compulsive alcohol-drinking behavior. There is increasing evidence that the dorsal striatum is a key hub in the regulation of cognitive flexibility (6, 7). We found that chronic alcohol consumption impaired the reversal of action-outcome contingency, indicating behavioral inflexibility. It is highly likely that this behavioral change is due to effects on the dorsal striatum (45, 46). Within this brain region, CINs play a critical role in regulating reversal learning (9, 10), which is essential in the reversal phase but not in the initial memory acquisition (9)—a fact that highlights the importance of CINs activity for new state formation, or the revision of behavior to accommodate a new situation (9, 47). Given that the present study found that chronic alcohol consumption and withdrawal affected glutamatergic transmission from the thalamus to striatal CINs, it is highly possible that this disruption induces a deficit in goal-directed action selection (9). This prediction was supported by our devaluation results, which indicated that chronic alcohol intake and withdrawal impaired devaluation during contingency reversal but did not impair contingency acquisition. Taken together, these findings suggest that chronic alcohol intake functionally impaired the flexibility of goal-directed behavior mediated by striatal circuits.

Having observed these behavioral effects of alcohol, we investigated whether the thalamic to CINs circuits were affected. Most previous *in vitro* and *in vivo* studies have demonstrated that thalamic stimulation produces burst-pause activity in CINs, which modulate D1-and D2-MSNs (17, 25). Thalamic glutamatergic transmission to CINs is, therefore a key component of this circuit(17). By selectively activating thalamic inputs to the DMS, the present study demonstrated that thalamic input modulated CIN activity and thus controlled the striatal MSN network. Furthermore, we found that chronic alcohol consumption decreased thalamic glutamatergic transmission to CINs. This result is consistent with the acute alcohol impairing the ability of thalamostriatal inputs to modulate a subsequent corticostriatal glutamatergic response in MSNs ^(3)^. Our study of paired-pulse ratios found that the probability of glutamate release did not decrease, indicating that this effect was not mediated by a decrease in thalamic activity (Fig. 2). A previous acute alcohol exposure study reported a decrease in evoked GABAergic MSN responses, with no change in the paired-pulse ratio (3). The mechanisms underlying this reduced thalamic input to DMS CINs need further investigation.

Interestingly, the present study employed a chronic alcohol consumption procedure, we found an increase in the spontaneous firing of CINs, in contrast to the inhibiting effect of acute alcohol on CINs firing (3). It is not uncommon that chronic and acute drugs have opposite effects due to the adaptation response of the organism. For example, acute morphine administration increased the spontaneous firing of dopamine neurons in the ventral tegmental area (VTA) (48, 49), while chronic morphine administration and withdrawal greatly reduced the spontaneous activity of VTA dopamine neurons (50, 51). We also found that acute alcohol suppressed the NMDA activity while chronic alcohol consumption enhanced NMDA function (52). As for the function of CINs it may not directly correlate with the baseline activity, in other words, increased baseline firing does not mean enhanced function of CINs. It has been shown the aged mice have increased CINs spontaneous firing and exhibit impairments in reversal learning of action-outcome contingency (10). Pharmacogenetic direct stimulation of CINs in the DMS did not alleviate the impairment of reversal learning in aged mice (10). It seems the extent that CINs can be modulated rather than their baseline firing plays a more important role in their function. With increased baseline firing, CINs could be less prone to be modulated, such as the shortened pause response observed in our study (Fig. 4). By selectively strengthening the thalamic inputs to DMS CINs, we were able to rescue the impairment of reversal learning in chronic alcohol consumption rats (Fig. 7).

CINs are a major source of acetylcholine within the striatum, and their dense terminals primarily synapse with MSNs. We generated triple transgenic mice in order to induce selective optogenetic excitation of CINs and to allow fluorescent identification of D1-MSNs. We found that direct optogenetic excitation of CINs elicited a stimulation-evoked firing response followed by a pause (Fig. 4). The burst-pause firing of CINs is intricately linked with dopamine activity in the striatum (53). Indeed, the pause duration was reduced by blocking dopamine D2Rs (Supplementary Fig. 4). This result is consistent with the finding that the thalamically-evoked pause is dependent upon dopamine release and DR activation(17). Therefore, direct optogenetic stimulation of CINs might exert a complex and powerful influence on specific types of striatal outputs. Cholinergic muscarinic M4 receptors, functionally coupled with the NMDAR, are only expressed in D1-MSNs(54). In addition, acetylcholine produced a prolonged enhancement of postsynaptic responsiveness in D2-MSNs. Our findings showed that the burst stimulation of CINs at 15 Hz, which is close to the burst firing frequency observed under physiological conditions (17), suppressed NMDAR-mediated glutamatergic inputs onto D1-MSNs and facilitated AMPAR-mediated glutamatergic transmission in D2-MSNs. The depression of D1-MSNs and facilitation of D2-MSNs by our direct optical activation of CINs was consistent with previous studies that employed electrical stimulation of the thalamus (17). The integrated effect on DMS MSNs, namely a decrease in the D1-direct pathway output and an increase in the D2-indirect pathway output, is to activate the striatopallidal network to suppress action (“No-Go”). Our results showed that chronic alcohol intake disrupted CIN-mediated depression of D1-MSNs and facilitation of D2-MSNs. A previous study also found that glutamatergic transmission increased in D1-MSNs after alcohol consumption (23). The effect of this disruption, which increases the relative activity of D1-MSNs and reduces that of D2-MSNs, is to reduce action suppression and make a “Go” outcome more likely. The discovery of these mechanisms provides a deep understanding of how alcohol consumption impacts thalamostriatal-CIN-MSN connectivity and thus promotes behavioral inflexibility.

In summary, DMS CINs modulate striatal circuits via burst-pause firing, which is triggered by inputs from the thalamus. Alcohol consumption disrupts this modulation by reducing thalamic excitation of CINs, and increasing spontaneous CIN activity. Our research demonstrated that alcohol attenuated the CIN-mediated inhibition of glutamatergic transmission in D1-MSNs and the CIN-mediated short-term facilitation of glutamatergic transmission in D2-MSNs. These effects have the potential to impair cognitive flexibility. Our findings provide an evidence base for the development of new therapeutic strategies to enhance cognitive flexibility in alcohol use disorder.

## Methods

### Animals

ChAT-eGFP, VGluT2-Cre, ChAT-Cre, and D1-tdT mice were purchased from the Jackson Laboratory. All mice were backcrossed onto a C57BL/6 background. VGluT2-Cre or ChAT-Cre mice were crossed with Ai32 to generate VGluT2-Cre;Ai32 or ChAT-Cre;Ai32 lines. VGluT2-Cre (or ChAT-Cre) and ChAT-eGFP (or D1-tdT) mice were crossed with Ai32 to generate VGluT2-Cre;Ai32;ChAT-eGFP or ChAT-Cre;Ai32; D1-tdT triple transgenic mice. Both male and female mice were used for electrophysiology studies. Male Long-Evans rats (3 months old) purchased from Harlan Laboratories were used for behavioral testing. Long Evans-Tg(ChAT-Cre) rats were purchased from Rat Resource & Research Center. Animals were housed individually at 23°C under a 12-h light:dark cycle, with lights on at 7:00 A.M. Food and water were provided ad *libitum.* All animal care and experimental procedures were approved by the Institutional Animal Care and Use Committee and were conducted in accordance with the National Research Council *Guide for the Care and Use of Laboratory Animals.*

### Reagents

PD 102807, dihydro-β-erythroidine hydrobromide, and DNQX (6,7-dinitroquinoxaline-2,3-dione) were purchased from Tocris. LY367385, mecamylamine hydrochloride, methyllycaconitine citrate, sulpiride, scopolamine, picrotoxin, and others were obtained from Sigma.

### Behavioral Procedures

#### Intermittent-access to 20% alcohol 2-bottle choice drinking procedure

This procedure was conducted as described previously (23, 24, 31–34, 55). Briefly, animals were given concurrent access to one bottle of alcohol (20%, in water) and one bottle of water for 24-h periods, which were separated by 24- or 48-h periods of alcohol deprivation. Alcohol intake (g/kg/day) was calculated by determining the weight of 20% alcohol solution consumed and multiplying this by 0.2. Water control animals only have access to water.

#### Magazine training

This procedure was adapted from Bradfield et al. (35). After 5 days of food restriction, rats were trained for magazine entries for 20 min on two consecutive days. During these training sessions, a reinforcer (either a food pellet or 0.1 mL sucrose solution) was delivered along with illumination of the magazine light for 1 sec with a random interval between each reinforcer (on average 60 sec). The house light was illuminated throughout the session, and no levers were available during magazine training. An equal number of rats received either 20 food pellets or 20 sucrose deliveries during the first training session and were then switched to receive the other reward in the second training session.

#### Acquisition of initial contingencies

Following magazine training, rats were trained to access different reinforcers via lever pressing over the next 10 days. Each session consisted of 4 blocks (2 blocks per lever), separated by a 2.5-min timeout during which no levers were available, and all lights were extinguished. Only one lever was available during each block (pseudorandom presentation), which lasted for 10 min or until 10 reinforcers had been earned. For half of the animals in each group, the left lever was associated with food pellet delivery and the right lever with sucrose solution delivery. The remaining animals were trained using the opposite pairs of action-outcome contingencies. Lever training started with a fixed ratio 1 (FR1) schedule in which every lever press resulted in the delivery of a reinforcer. After 2 days of FR1 training, the training schedule was elevated to a random ratio 5 (RR5) schedule for the next 3 days, during which a reinforcer was delivered after an average of 5 lever presses. An RR10 training schedule was then employed for 3 days, followed by an RR20 schedule for the final 2 days.

#### Devaluation test

After the final RR20 training, devaluation testing was performed for 2 days. On both days, rats were habituated in a dark, quiet room (different from the operant room) for 30 min, then were given *ad libitum* access to either the food pellets (25 g placed in a bowl) or the sucrose solution (100 mL in a drinking bottle) in a devaluation cage for 1 h. The devaluation cage was similar to their home cage but did not contain bedding. The rats were then placed in the operant chamber for a 10-min extinction choice test. Both levers were extended during this test, but no outcomes were delivered in response to any lever press. On the second devaluation day, the rats were pre-fed, as described, with the other reward before repeating the same extinction test. Lever presses (LP) were recorded, and those on the lever that the rat had learned to associate with the non-devalued reward were termed LP_valued_, while those on the lever associated with the devalued reward were termed LP_devalued_. The devaluation index [(LP_valued_ - LP_devalued_)/(LP_valued_ + LP_devaluted_)] was then used to determine the extent of goal-directed versus habitual behavior.

#### Contingency reversal and devaluation testing

After the devaluation test, rats were retrained on their current action-outcome contingencies for 1 day. The contingencies were then reversed so that the lever that previously delivered food now delivered sucrose, and the rats were trained using the RR20 schedule. All other procedures were unchanged. The contingency reversal training lasted for 4 days. The rats then underwent devaluation testing again using the procedure described above.

### Electrophysiology

Slice electrophysiology was performed as previously described (24, 55). Animals were sacrificed 24 h after their last alcohol (or control water) consumption, and 250-μm coronal sections containing the striatum were prepared in an ice-cold cutting solution containing (in mM): 40 NaCl, 148.5 sucrose, 4 KCl, 1.25 NaH_2_PO_4_, 25 NaHCO_3_, 0.5 CaCl_2_, 7 MgCl_2_, 10 glucose, 1 sodium ascorbate, 3 sodium pyruvate, and 3 myoinositol, saturated with 95% O_2_ and 5% CO_2_. Slices were then incubated in a 1:1 mixture of cutting solution and external solution at 32°C for 45 min. The external solution contained the following (in mM): 125 NaCl, 4.5 KCl, 2.5 CaCl_2_, 1.3 MgCl_2_, 1.25 NaH2PO4, 25 NaHCO3, 15 sucrose, and 15 glucose, saturated with 95% O_2_ and 5% CO_2_. Slices were then maintained in external solution at room temperature until use.

Slices were perfused with the external solution at a flow rate of 3-4 mL/min at 32°C. The CINs and MSNs in the DMS were identified either by differential interference contrast or by fluorescence. Whole-cell patch-clamp and cell-attached recordings were made using a MultiClamp 700B amplifier controlled by pClamp 10.4 software (Molecular Devices). For cell-attached and whole-cell current-clamp recordings, we used a K^+^-based intracellular solution containing (in mM): 123 potassium gluconate, 10 HEPES, 0.2 EGTA, 8 NaCl, 2 MgATP, 0.3 NaGTP (pH 7.3), with an osmolarity of 270–280 mOsm. For whole-cell voltage-clamp recordings, we used a Cs-based solution, containing (in mM): 119 CsMeSO4, 8 TEA.Cl, 15 HEPES, 0.6 EGTA, 0.3 Na_3_GTP, 4 MgATP, 5 QX-314.Cl, 7 phosphocreatine. The pH was adjusted to 7.3 with CsOH.

For measurement of spontaneous CIN firing, cell-attached recordings were conducted in the voltage-clamp mode. In whole-cell current-clamp recordings, evoked action potentials were elicited by 500-ms stepped current injections at 30-pA increments from −120 pA to +120 pA. Optogenetically-evoked CIN firing was induced by light stimulation (473 nm, 2 ms, 15 Hz, 10 pulses) through the objective lens. Bipolar stimulating electrodes were positioned 100-150 μm away from the recording electrode that was used to record glutamatergic transmission in MSNs. To measure NMDAR-EPSCs, the neurons were recorded in the presence of DNQX and with magnesium-free external solution. All of the measurements were conducted in the presence of the GABA_A_ receptor antagonist, picrotoxin (100 μM). The experiments in Figure 5 were conducted in the presence of the mGluR1/5 antagonists, LY367385 (10 μM).

### Stereotaxic surgery and Histology

The rabies helper viruses (AAV8-DIO-RG and AAV8-DIO-TVA-mCherry), AAV-Chrimson-tdT, AAV-FLEX-Chrimson-tdT, and AAV-Chronos-GFP were purchased from the University of North Carolina Vector Core. The pseudotyped rabies viruses, EnvA-SADΔG-mCherry and EnvA-SADΔG-GFP (2.04 ×10^8^ TU/mL), were obtained from the Salk Institute Vector Core.

Stereotaxic viral infusions were performed as described previously (23, 24, 28, 56). Briefly, mice were anesthetized using isoflurane and mounted in a rodent stereotaxic frame (Kopf). The skin was opened to uncover the skull and expose Bregma and Lambda, and the location of the desired injection site. A three-axis micromanipulator was used to measure the spatial coordinates for Bregma and Lambda. Small drill holes were made in the skull at the appropriate coordinates, according to the Paxinos atlas (57). Two microinjectors were loaded with 0.5 μL of a 1:1 mixture of AAV8-DIO-RG and AAV8-DIO-TVA-mCherry, and then lowered into the pDMS (AP: 0.0 mm, ML: ± 1.87 mm, DV: −2.90 mm). This helper virus mixture was infused into the brain at a rate of 0.1 μL/min. To avoid backflow of the virus, microinjectors were left in place for 10 min after the infusion was complete. Following their removal, the skin was sutured and the mice were allowed to recover for 3 weeks prior to the infusion of pseudotyped rabies virus (EnvA-SADΔG-mCherry or EnvA-SADΔG-eGFP). The rabies virus was injected at the same site and using the same injection volume as the initial helper virus injection. To prevent coincident rabies infection along the injection tract, the rabies virus was infused into adapted coordinates (AP, 0.0 mm; ML, ± 2.42 mm; DV, −2.94 mm) at an angle of 10 degrees (58) to the previous injection. The modified coordinates were calculated by measuring from the midline and parallel to the dorsal-ventral axis. The coordinates for mice AAV-Chrimson-tdT (0.5 μL) PfN injection were AP, −2.2 mm; ML, ± 0.7 mm; and DV, −3.5 mm. ChAT-Cre rats, DMS (AAV-FLEX-Chrimson-tdT): AP, 0.0 mm; ML, ± 2.8 mm; and DV, −4.85 mm; PfN (AAV-Chronos-GFP): AP, −4.2 mm; ML, ± 1.25 mm; and DV, −6.2 mm. For rats, 1 μl to 1.2 μl of the virus was infused in each hemisphere. After virus injections, bilateral optical fiber implants (300-μm core fiber secured to a 1.25-cm ceramic ferrule with 5 mm of fiber extending past the end of the ferrule) were lowered into the DMS right on the top of virus injection sites. Coordinates: AP, 0.0 mm; ML, ± 2.8 mm; and DV, −4.8 mm. Implants were secured on the skull using metal screws and dental cement (Henry Schein) and covered with denture acrylic (Lang Dental). The incision was closed around the head cap and the skin vet-bonded to the head cap. Rats were monitored for 1 week or until they resumed normal activity.

The histology procedure was performed as described previously (24, 56). Briefly, mice were anesthetized and perfused intracardially with 4% paraformaldehyde in phosphate-buffered saline (PBS). Whole brains were taken out and placed into 4% paraformaldehyde in PBS for post-fixation overnight (4°C), then moved to 30% sucrose in PBS (4°C) and allowed to sink to the bottom of the container before preparing for sectioning. Frozen brains were cut into 50-μm coronal sections on a cryostat. A confocal laser-scanning microscope (Fluorview-1200, Olympus) was used to image these sections with a 470-nm laser (to excite eYFP and GFP) and a 593-nm laser (to excite tdT). All images were processed using Imaris 8.3.1 (Bitplane, Zurich, Switzerland).

#### Statistical analysis

All data are expressed as the mean ± SEM. Statistical significance was assessed using the unpaired or paired *t* test or two-way RM ANOVA followed by the *Tukey* test for *post hoc* comparisons. Statistical significance was set at *p* < 0.05.

## Author contributions

J.W. conceived, designed, and supervised all the experiments in the study. T.M. and Z.H. contribute equally to this research. The order of co-first author is determined by who completed the first draft of the manuscript. T.M. wrote the first draft of the manuscript and J.W., T.M., Z.H., Y.C., L.S., R.S., and Y.Z revised the manuscript. Z.H., T.M., and X.Z. designed and performed electrophysiology experiments and analyzed the data. Z.H., X.X., and M.C. designed and performed the behavior experiments and analyzed the data. H.G. and X.W. conducted histology experiments.

## Acknowledgements

We appreciate Drs. David Lovinger’s and David Earnest’s critical comments on our manuscript. We thank Sebastian Melo and Jared Jarger for technical assistance. This research was supported by NIAAA U01AA025932 (J.W.), R01AA021505 (J.W.), and R01AA027768.

The authors report no biomedical financial interests or potential conflicts of interest.

