## Supplementary Figures 1-5 for "Chronic alcohol drinking persistently suppresses thalamostriatal excitation of cholinergic neurons to impair cognitive flexibility"

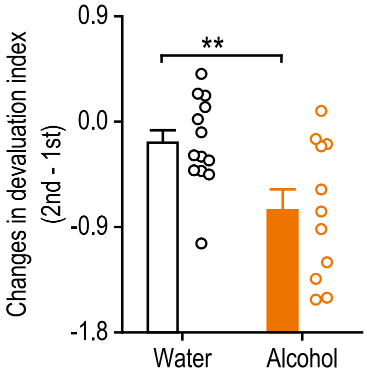


**Supplementary Figure 1.** Chronic alcohol-drinking rats shift their action strategy from goal-directed during initial A-O acquisition to habitual during reversed A-O learning. The difference between the 1$^{st}$ and 2$^{nd}$ devaluation index represents the degree of change in their action strategy. Black dots represent individual changes in the water group, and red dots represent individual changes in the alcohol group; $^{**}p < 0.01$ by unpaired *t*-test, n = 13 rats (Water) and 11 rats (Alcohol).

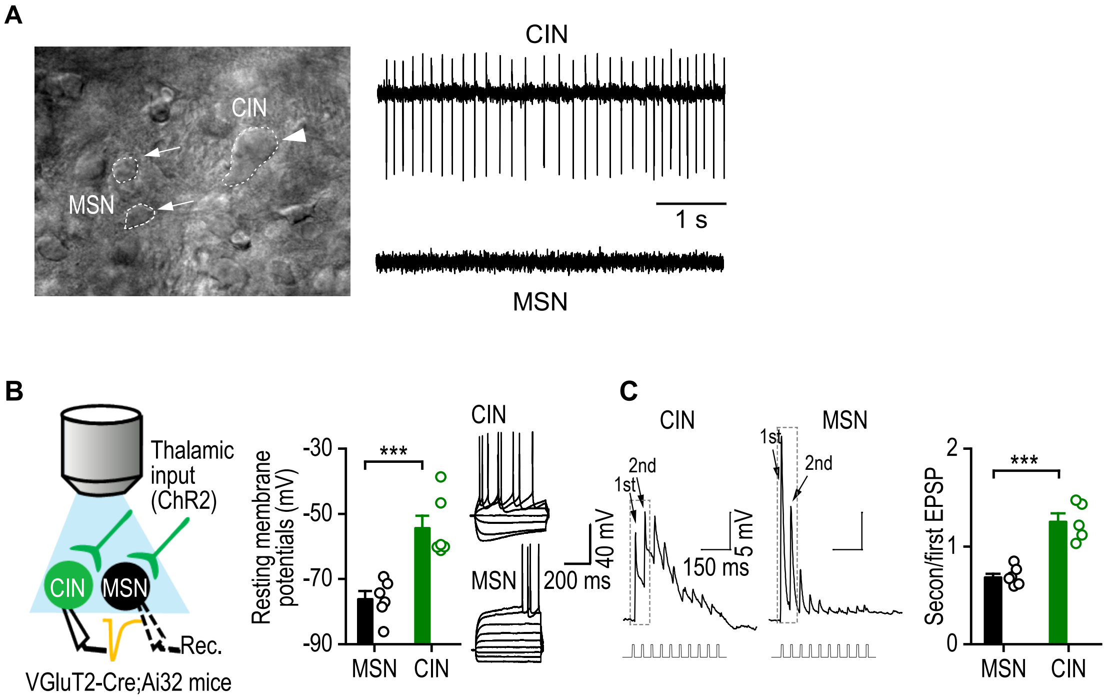


**Supplementary Figure 2.** The distinct features of CINs and MSNs in the DMS. **(A)** DIC image showing a CIN (arrowhead) and MSNs (arrows), and sample traces of spontaneous firing in CINs (top), but not in MSNs (bottom) using cell-attached recording. **(B)** Schematic diagram showing light stimulation of ChR2-expressing thalamic inputs and whole-cell recording of CINs or MSNs in DMS slices prepared from VGluT2-Cre;Ai32 or VGluT2-Cre;Ai32;ChAT-eGFP mice. Characterization of CINs and MSNs using their distinct resting membrane potentials and responses to current injection. Left, CINs had higher resting membrane potentials than did MSNs; ***$p < 0.001$ by unpaired *t* test, $n = 6$ neurons from 3 mice (6, 3) per group. Right, less current was needed to generate action potentials in CINs than in MSNs using 500-ms step-current injections, starting at -120 pA with 30-pA increments. **(C)** Left and middle, sample traces showing that a train of light stimulation (2 ms, 10 pulses, 15 Hz) of thalamic inputs elicited facilitation of the second EPSP in CINs, but the depression of the second EPSP in MSNs. Right, the ratio of the second:first EPSP was significantly higher in CINs than in MSNs; ***$p < 0.001$ by unpaired *t* test, $n = 5, 4$ (CINs) and 7, 4 (MSNs).

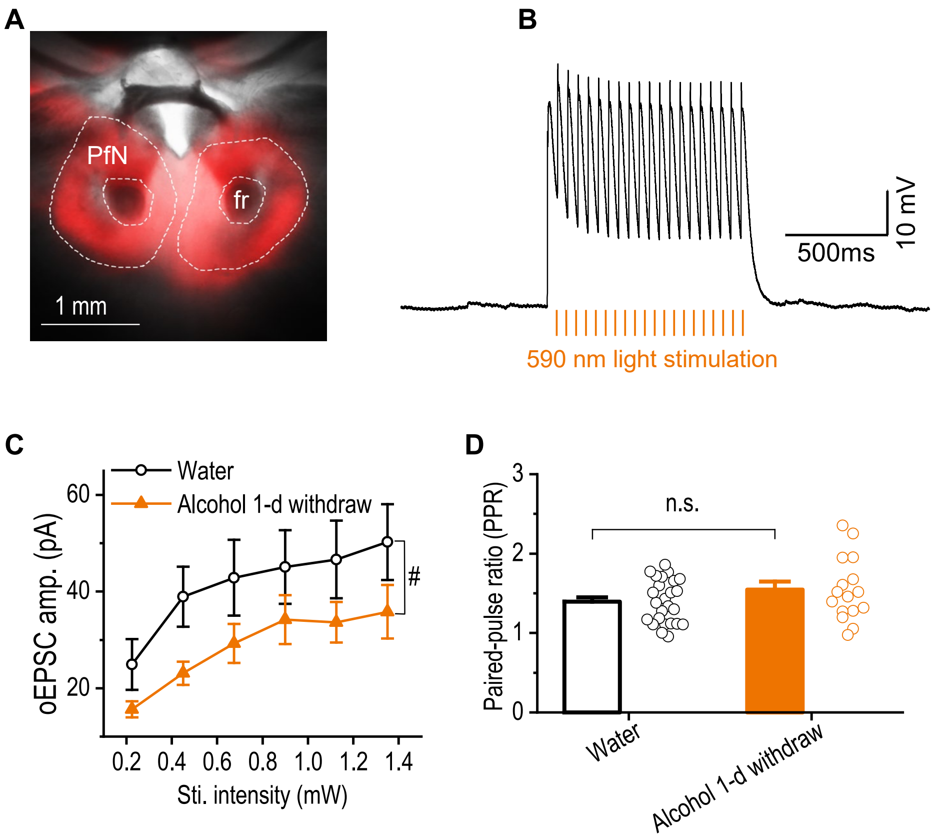


**Supplementary Figure 3.** Chronic alcohol intake reduces thalamic excitatory inputs onto DMS CINs of mice injected with AAV-Chrimson-tdT**. (A)** Optical image showing the expression of AAV-Chrimson-tdT in the PfN of the thalamus. **(B)** A representative trace showing the activation of a neuron expressing Chrimson by optical stimulation in the PfN. **(C)** Input-output curves of oEPSC amplitudes in CINs from mice injected with AAV-Chrimson-tdT and exposed to alcohol or water; $^{\#}p < 0.05$ by two-way RM ANOVA; $n = 12$, 2 (Water) and 12, 2 (Alcohol). **(D)** Paired-pulse ratios in mice injected with AAV-Chrimson-tdT and exposed to alcohol or water; $p > 0.05$ by unpaired $t$ test, $n = 27$, 5 (Water) and 16 2 (Alcohol).

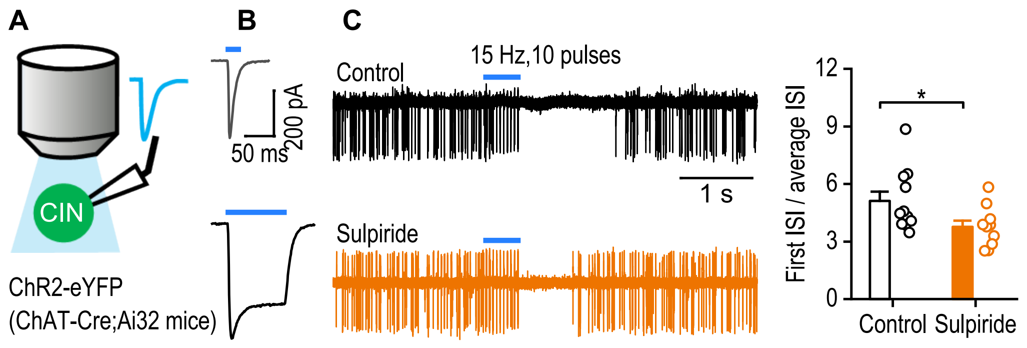


**Supplementary Figure 4.** Direct optical stimulation of CINs generates a pause mimicking the thalamostriatally-evoked burst-pause response**. (A)** Diagram showing light stimulation of, and whole-cell recording in, ChR2-expressing DMS CINs from ChAT-Cre;Ai32 mice. **(B)** Short (2 ms) or prolonged (100 ms) light stimulation of DMS CINs elicited a short-lived (top) or sustained (bottom) depolarization. **(C)** Inhibition of D2Rs reduced the pause response of DMS CINs to direct light excitation. Left, superimposed sample traces show that light-mediated CIN stimulation induced a "burst-pause" response in the absence (top) and presence (bottom) of the D2R antagonist, sulpiride (20 μM). Right, the ratio of the first inter-spike interval (ISI) after light stimulation was divided by the average ISI measured before stimulation (500 ms). This ratio was lower in the sulpiride group than in the control group; $^*p < 0.05$ by unpaired $t$ test, $n$ = 11,4 (Control) and 10,4 (Sulpiride).

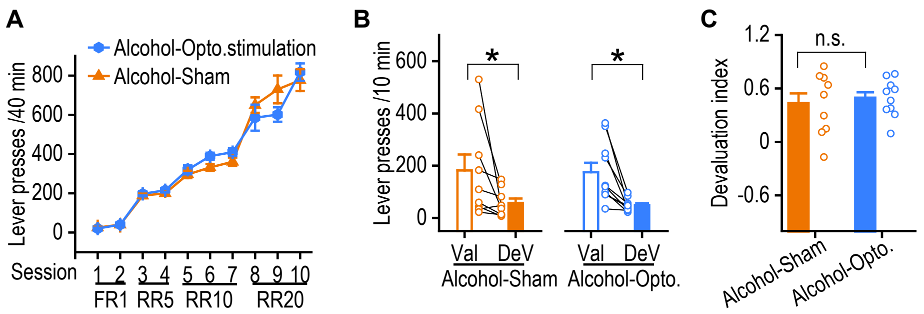


**Supplementary Figure 5.** Group performance for the initial learning before light stimulation**. (A)** Two groups showed no significant difference in total lever presses during the acquisition of the initial contingencies, moving from a fixed ratio 1 (FR1) schedule to random ratio 5 (RR5), RR10, and RR20 schedules as indicated; two-way RM ANOVA, n = 9 rats (Alcohol-Sham) and 10 rats (Alcohol-Opto). **(B)** Outcome-specific devaluation testing showed that both sham and light stimulation groups pressed the DeV lever significantly fewer times than the Val lever; $^*p < 0.05$ by paired $t$ test, n = 9 rats (Alcohol-Sham) and 10 rats (Alcohol-Opto.stimulation). **(C)** The devaluation index, defined as (Val - DeV)/(Val + DeV), did not differ significantly between the two groups; n.s., not significant by unpaired $t$ test, n = 9 rats (Alcohol-Sham) and 10 rats (Alcohol-Opto).
